# TRANSCUP: a scalable workflow for predicting cancer of unknown primary based on next-generation transcriptome profiling

**DOI:** 10.1101/774315

**Authors:** Peng Li

## Abstract

**Summary:** Cancer of unknown primary site (CUP) accounts for 5% of all cancer diagnoses. These patients may benefit from more precise treatment when primary cancer site was identified. Advances in high-throughput sequencing have enabled cost-effective sequencing the transcriptome for clinical application. Here, we present a free, scalable and extendable software for CUP predication called TRANSCUP, which enables (1) raw data processing, (2) read mapping, (3) quality re-port, (4) gene expression quantification, (5) random forest machine learning model building for cancer type classification. TRANSCUP achieved high accuracy, sensitivity and specificity for tumor type classification based on external RNA-seq datasets. It has potential for broad clinical application for solving the CUP problem.

**Availability:** TRANSCUP is open-source and freely available at https://github.com/plsysu/TRANSCUP

## Introduction

The Cancer of unknown primary site (CUP) is a heterogeneous group of cancers in the absence of an identifiable primary tumor despite a standard diagnostic approach and accounts for approximately 5% of all cancer diagnoses. The biologic mechanisms underlying this phenomenon is unknown. CUP patients can be given site specific therapy with significant improvement in clinical out-come compared with empirical chemotherapy when cancer primary site was identified (Pavlidis and Pentheroudakis, 2012; Varadhachary and Raber, 2014).

Currently, many molecular diagnostic methods have been widely applied in clinic, including RT-PCR (Overman, et al., 2016), microRNA RT-PCR (Rosenfeld, et al., 2008), gene expression micro-array (Pillai, et al., 2011) and DNA methylation microarray (Moran, et al., 2016). Advances in high-throughput sequencing have enabled cost-effective sequencing the transcriptome for clinical application. RNA-Seq based predication algorithm have been proposed using TCGA’s RNA-Seq RSEM expression value and validated on both RNA-Seq and microarray dataset (Flynn, et al., 2018). How-ever, data analysis pipelines from raw FASTQ data to final tumor type predication are currently not fully implemented and validated computationally.

Here, we present a free, scalable and extendable software for CUP predication called TRANSCUP, which comprises modules for raw data processing, read mapping, quality report, gene expression quantification and building a random forest model for cancer type classification. It achieved high accuracy, sensitivity and specificity for tumor type classification based on external RNA-seq datasets. It has potential for broad clinical application for solving the CUP problem.

## Methods

The TRANSCUP workflow is illustrated in Figure 1. To build CUP classifier, we use TCGA RNA-Seq data as training dataset, followed by sample and feature selection, data transformation, data normalization and random forest model building. To predict new tumor samples, we build a bioinformatics pipeline to process data from raw FASTQ reads to tumor type predication result. The de-tails for implementation are the followings:

**Fig. 1.**
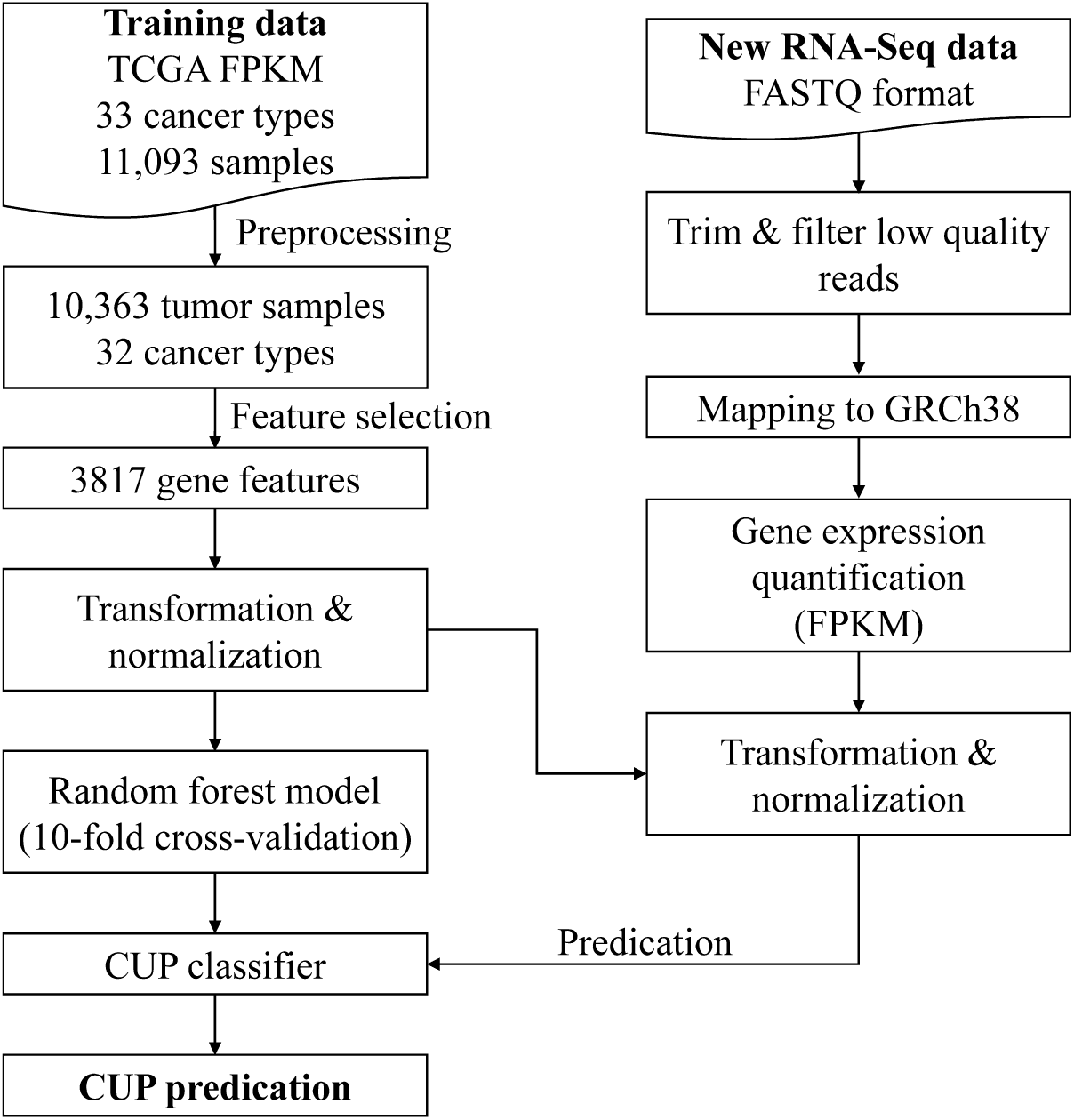
Workflow of TRANSCUP.

### Training data source

TCGA harmonized gene expression quantification data and clinical data were retrieved from GDC portal (https://portal.gdc.cancer.gov/) via TCGAbiolinks (Colaprico, et al., 2016). Totally, 32 cancer types (COAD and READ were combined to CRC) and 10,363 tumor samples were used for model building.

### Preprocess and feature selection

All FPKM expression data was log2-transformed, and only genes with (a) maximum log2 expression greater than 2.5 and (b) variance in log2 expression greater than 0.1 were retained. After filtering, the genes in each dataset was scaled to zero mean expression and unit variance. The mean and standard variance learned from training dataset for each gene were used to normalize the new data. Feature genes were selected by two criterion: 1) those genes were different expressed by one tumor type to other tumor types; 2) those genes were expressed more higher in one tumor type than other tumor types, which evaluated by t-test and log2-fold change, respectively. The top 100, 150, 200 genes’ log2 fold change value or log2 fold change value larger than 2.5 or 3 were compared to find the best feature selection method.

### Random forest model

The random forest algorithm was used for machine learning model building. To avoid overfitting, model was validated by 10-fold cross-validation. The final model was trained on all training data with the best hyper-parameters.

### Bioinformatics pipeline

The raw reads were cleaned prior to following analysis by Trimmomatic (Bolger, et al., 2014). Clean reads were mapped to the human genome GRCh38 by STAR (Dobin, et al., 2013) using the 2-pass model. Gene read counts were calculated using HTSeq-count (Anders, et al., 2015). GENCODE v22 GTF file was used to alignment and HTSeq-count. To quantify gene expression, the fragments per kilobase of transcript per million mapped reads (FPKM) values of each gene were calculated. To evaluate the RNA-seq data quality, multiple metrics include yield, alignment, GC bias, rRNA content, regions of alignment (exon, intron and intragenic), continuity of coverage, 3_′_/5_′_ bias and count of detectable genes, among others were calculated by RNA-SeQC (DeLuca, et al., 2012).

### Snakemake workflow

To make this software extendable and scalable, we adopted snakemake (Koster and Rahmann, 2012; Singer, et al., 2018) workflow engine to chain the tools, databases, and config files together. This enables users to easily make use of any cluster environment for processing and conveniently manage tools, databases and pipelines.

## Results

We found that TOP200 was the best feature selection method after compared with other 4 methods mentioned above (Supplemental Table S1). 3817 genes were retained as feature genes (Supplemental Table S2). In the training phase, we performed 10-fold cross-validation on all training data to avoid overfitting and find best hyper-parameters. The accuracy, kappa, standard deviation of accuracy and standard deviation of kappa were 0.961, 0.959, 0.003292 and 0.003457, respectively. The median of sensitivity and specificity across 32 cancer types were 0.969 and 0.999, respectively (Supplemental Table S3). We used five external public RNA-seq datasets (Supplemental Table S4) to evaluate the performance of TRANSCUP. These datasets totally contain 557 samples and across 4 different cancer types including OSCC (HNSC), CRC, lung cancer (LUAD/LUSC) and BRCA. Notably, the overall accuracy of TRANSCUP was 98.7%, totally 7 samples were misclassified (Supplemental Table S5).

## Conclusions

In this article, we described the TRANSCUP package for tumor type predication. TRANSCUP has been validated using 557 external samples and was more accurate than other methods for cancer type classification. Furthermore, TRANSCUP can analyze data from raw FASTQ to final cancer type predication results, and is more scalable and extendable than other methods. Users can train other kinds of models like deep learning models to extend its capability. However, this package needs to be validated on more external RNA-Seq datasets which including more diverse cancer types when data are available. Its actual clinical effect has to be verified by further experiments and clinical trials in the future.

