## Supplemental Table S4 for "TRANSCUP: a scalable workflow for predicting cancer of unknown primary based on next-generation transcriptome profiling"

**Supplemental Table S4: The source of external public RNA-seq datasets**

| **Cancer type** | **No. of samples** | **Reference** | **Accession number** |
| --- | --- | --- | --- |
| OSCC(HNSC) | 39 | (Chen, et al., 2017) | SRP078156 |
| CRC | 10 | (Lee, et al., 2016) | SRR2089755 |
| LUAD/LUSC | 45 | (Jia, et al., 2018) | GSE112996 |
| BRCA | 357 | (Jiang, et al., 2019) | OEP000155 |
| CRC | 106 | (Vasaikar, et al., 2019) | PRJNA514017 |

Chen, T.-W.*, et al.* APOBEC3A is an oral cancer prognostic biomarker in Taiwanese carriers of an APOBEC deletion polymorphism. *Nature Communications* 2017;8(1):465.

Jia, Q.*, et al.* Local mutational diversity drives intratumoral immune heterogeneity in non-small cell lung cancer. *Nature Communications* 2018;9(1):5361.

Jiang, Y.Z.*, et al.* Genomic and Transcriptomic Landscape of Triple-Negative Breast Cancers: Subtypes and Treatment Strategies. *Cancer Cell* 2019;35(3):428-440 e425.

Lee, J.-R.*, et al.* Transcriptome analysis of paired primary colorectal carcinoma and liver metastases reveals fusion transcripts and similar gene expression profiles in primary carcinoma and liver metastases. *BMC Cancer* 2016;16(1):539.

Vasaikar, S.*, et al.* Proteogenomic Analysis of Human Colon Cancer Reveals New Therapeutic Opportunities. *Cell* 2019;177(4):1035-1049 e1019.
